# High-resolution Dynamic Human Brain Neural Activity Recording Using 3T MRI

**DOI:** 10.1101/2023.05.31.542967

**Authors:** Yifei Zhang, Kaibao Sun, Jianxun Ren, Qingyu Hu, Yezhe Wang, Shiyi Li, Tienzheng Chen, Na Xu, Ning Guo, Xiaoxuan Fu, Xuan Liu, Zhenen Cao, Jia-hong Gao, Hesheng Liu

## Abstract

Despite extensive research over decades, a non-invasive technique capable of capturing neural activities in the human brain with high spatiotemporal resolution is still lacking. The recently proposed direct imaging of neuronal activity (DIANA) using ultrahigh-field magnetic resonance imaging (MRI) has shown promise, but the translation from anesthetized mice to awake humans poses a significant challenge. Here we present Time Resolved ImaGing of Global Electroneurophysiological Record (TRIGGER), a novel technique that enables the direct detection of neural activity in the awake human brain using 3T MRI. In 18 participants, visual responses were captured at 5-mm spatial resolution and 1.4-ms temporal resolution. Importantly, the delay in stimulus presentation reliably corresponded to the latency of neural responses on a millisecond scale. Furthermore, when stimuli were presented to one visual field, the responses in two hemispheres exhibited the expected time difference. This non-invasive mapping approach holds the potential to elucidate neural dynamics underlying human brain function and disorders.

## Introduction

Breakthroughs in cognitive neuroscience are often driven by advances in recording techniques of neural activity. Historically, extensive endeavors have been made to record neural activities in vivo at high resolutions (*1*). In animal studies, the single-unit recording (*2*), patch clamp technique (*3*), Calcium imaging (*4*), and two-photon microscopy (*5*) have been widely used, offering high spatiotemporal resolution, but their invasive nature and limited recording scale and/or depth hindered the translation to awake humans for recording neural activities in large-scale functional networks (*6*). In noninvasive human studies, electroencephalography (EEG) and magnetoencephalography (MEG) have high temporal resolutions but suffer from limited spatial specificities (*7*). Conversely, magnetic resonance imaging (MRI), particularly the blood oxygenation level-dependent functional MRI (BOLD-fMRI), enables non-invasive whole-brain functional scanning, including deep nuclei, with high spatial resolution. Consequently, this neuroimaging technique has gained significant popularity over the past three decades, yielding numerous novel insights into brain organization, such as the identification of the fusiform face area (*8*) and the recently proposed somato-cognitive action network (*9*). However, the temporal resolution of BOLD-fMRI is constrained by the sluggishness of hemodynamic responses (*10, 11*). Therefore, there is still a missing piece in the puzzle of noninvasive technique for large-scale neural activity mapping in vivo that simultaneously possesses both high temporal and spatial resolution in awake human.

Recently, the direct imaging of neuronal activity (DIANA) with superior spatiotemporal resolution has been proposed and tested on anesthetized mice at 9.4T (*12*). The DIANA responses showed a high correlation with electroneurophysiological activity, making it a promising tool for measuring neural activity directly using MRI. Nevertheless, recent studies failed to obtain reliable DIANA signals in anesthesized mice at 15.2T (*13*) and in awake humans at 7T (*14*), likely due to the limited signal to noise ratio (SNR) of the MRI sequence. Translation from anesthetized mice to awake human hence remains a challenging task.

In order to tackle this challenge, we report here Time Resolved ImaGing of Global Electroneurophysiological Record (TRIGGER), an optimized MRI scanning method designed to achieve a high signal-to-noise ratio (SNR) and effectively record neural activities at a resolution of millimeters and milliseconds in awake human subjects.

## Results

To obtain high-resolution signals in awake humans within a reasonable acquisition time, we employed a two-dimensional (2D) imaging sequence that incorporated both frequency and phase encodings in a gradient echo train in a 3.0T MRI scanner (*15*). Each echo within an echo train captures a single line of a 2D k-space, and multiple repeated echo trains, synchronized with cyclic stimuli, capture a series of fully sampled 2D k-spaces. (Fig. 1A). By maintaining a short interval between two echoes, a high temporal resolution of 1.4 ms was achieved, ensuring precise and detailed temporal information. In comparison with the DIANA sequence, our sequence, with a relatively long repetition time (TR, 300 ms) and a large flip angle (FA, 40°), demonstrated improved SNR both theoretically and empirically within a time window of about 60 ms (Fig. 1B). The experimental data confirmed the SNR patterns that were estimated through numerical simulations for both sequences. While the SNR of TRIGGER exponentially decayed due to the free-induction decay (FID) of the gradient echo train, as opposed to the stable SNR of DIANA, the SNR of TRIGGER can be up to eight times higher than that of DIANA, providing an opportunity for better capturing weak neural signals. Importantly, to ensure precise synchronization between the presentation of stimuli and the acquisition of MRI data, we implemented a stimuli presentation system with minimal and stable latency (<1ms) through direct connection with the MRI spectrometer (Fig. 1C).

**Fig. 1.**
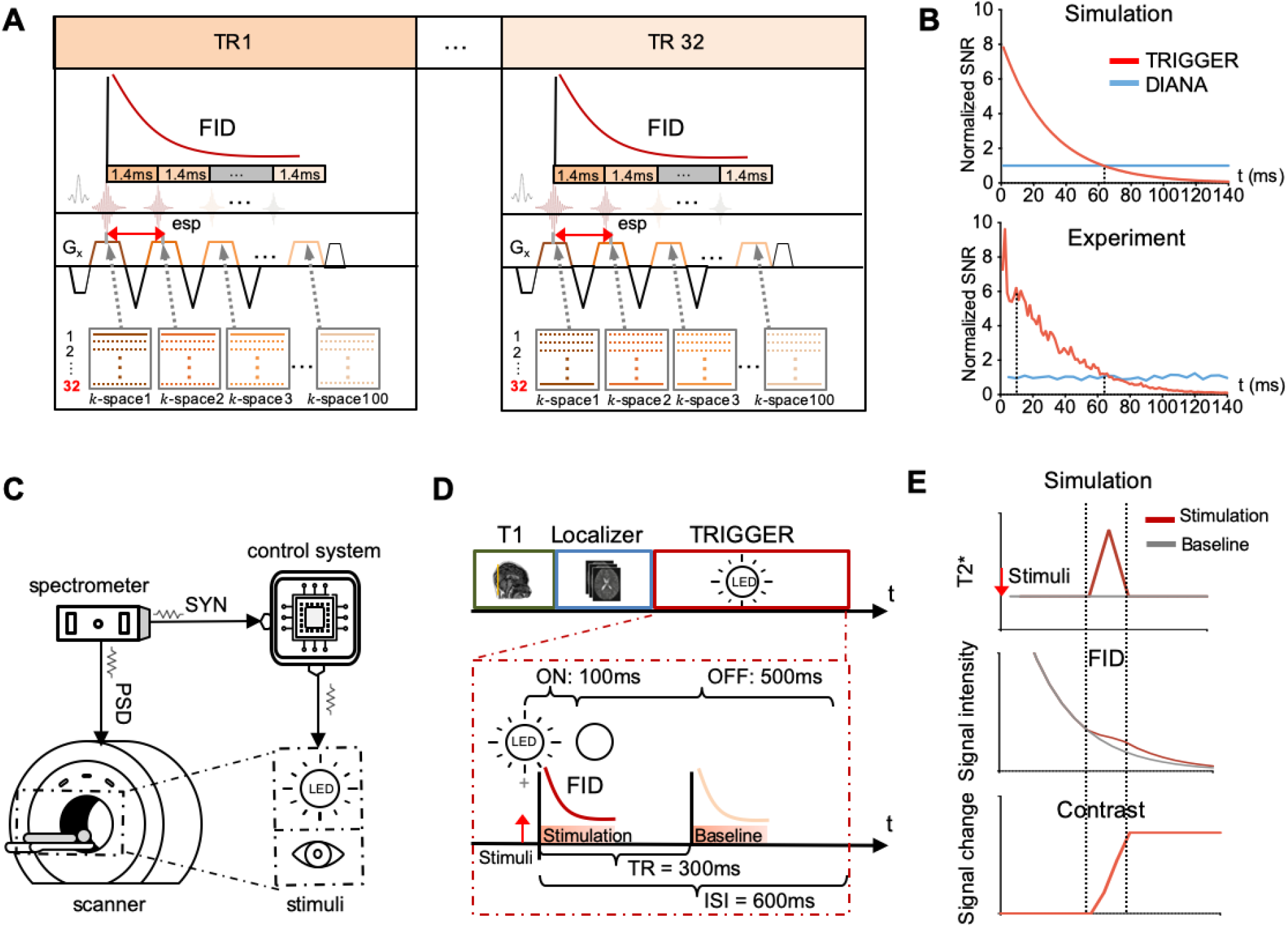
The TRIGGER sequence with a high SNR and the experimental design. (**A**) The TRIGGER approach used a 2D gradient-echo-train-based MRI imaging sequence with a 1.4 ms temporal resolution, determined by the time of echo spacing (esp). (**B**) The numerical simulation and empirical data reveal a greater SNR of TRIGGER compared to DIANA in the first 60 ms from the acquisition onset. (**C**) A low-latency stimulus presentation system was established through a microcontroller which receives the synchronizing pulse signals from the MR spectrometer and controls the stimuli delivery. PSD: pulse sequence diagram, LED: light-emitting diode. (**D**) In all subsequent experiments, T1w brain images and BOLD-fMRI localizer for early visual cortices were acquired to select the TRIGGER scanning slice. Multiple runs of TRIGGER scans were performed within each session, generating stimulation and baseline Free Induction Decay (FID) signals for each voxel in the selected slice. FID: free induction decay. (**E**) Illustration of data analysis steps. The neural activity may induce changes in T2* and thus yield different FID signals for stimulation condition compared to baseline. Signal changes represent the percentage contrast between FID signals of stimulation condition and baseline, reflecting cumulative changes in T2*.

To evaluate the capability of the TRIGGER approach in recording neural responses with high resolution, we performed an experiment in humans to activate the early visual cortices using flickering lights. Prior to the TRIGGER scans, we obtained T1-weighted brain images and performed a BOLD-fMRI visual localizer experiment to localize the early visual cortices. This allowed us to assess the effectiveness of the method in capturing neural activity in these specific brain regions. During the TRIGGER scans, flickering light stimuli were delivered to the upper visual field (light duration= 100 ms ; interstimulus interval (ISI) 600 ms, view angle = 3.4°), while participants maintained fixation at the center of the visual field (Fig. 1D). Each stimulus cycle was accompanied by the acquisition of two echo trains, each consisting of 100 echoes and lasting 140 ms. The first echo train was designed to capture the responses to visual stimulation and the second echo train captured the baseline activity (Fig. 1D). Changes in MRI signal induced by neural activities could be estimated by contrasting the percent signal change after the stimulation compared to the baseline recording. Our simulation indicated that changes in T2* will lead to an incremental MRI signal change in a stepwise manner (Fig. 1E).

We first conducted a pilot experiment with a single participant who underwent a 40-minute TRIGGER scanning session comprising 120 trials. Statistically significant TRIGGER responses were observed in the early visual cortex approximately 40 ms after stimulus onset (permutation test, z-statistic > 1.96, Fig. 2A). To test wether the TRIGGER responses capture neural activity at the millisecond scale, we employed a visual stimuli presentation paradigm with varying onset times. This experimental design allowed us to investigate the effects of different stimuli onset timings on the subsequent neural responses. In the first condition stimuli were presented 50 ms prior to the scanning (stimuli onset 1). In the other two conditions the visual stimuli onset were delayed by 10 ms (stimuli onset 2) and 20 ms (stimuli onset 3). The participant performed six runs (15 trials per run) under each condition, with a total scanning time of 1.5 hours. Signal changes were averaged across voxels within the visual region of interest (ROI) localized by the BOLD-fMRI task and across runs within the same condition. The MRI signal changes observed across the three conditions exhibited comparable dynamic patterns, but with noticeable temporal shifts (Fig. 2B). All three curves displayed gradual increases characterized by two distinct peaks. The latencies of these peaks, occurring approximately at 80 ms and 100 ms after stimulus onset, were found to be similar. These findings provide preliminary evidence of the high spatiotemporal resolution achieved through the implementation of the TRIGGER approach.

**Fig. 2.**
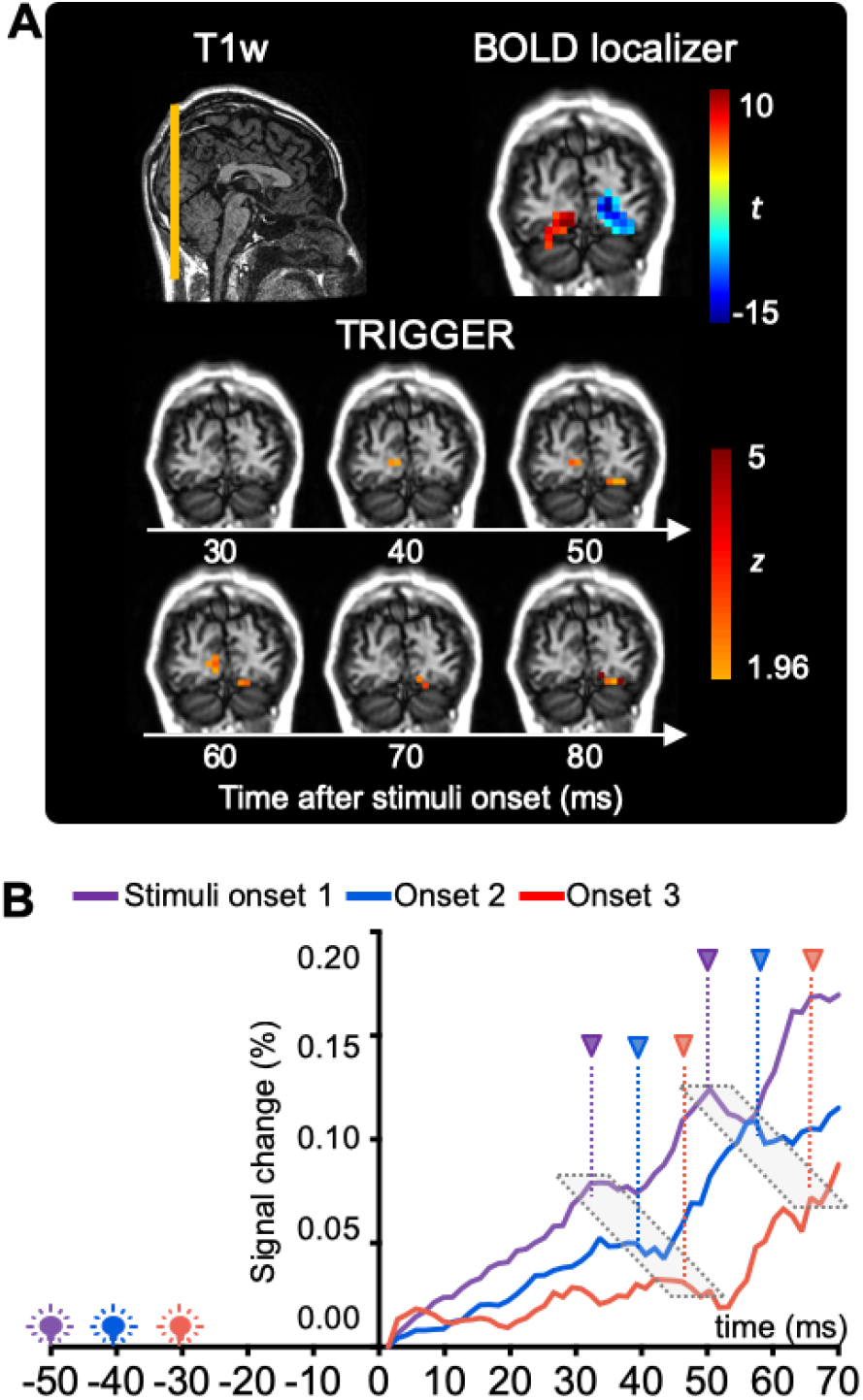
The TRIGGER responses to visual stimulus with high spatial and temporal resolution in an individual. **(A)** TRIGGER data were collected from a single human participant. The axial T1w image is illustrated for the coronial slice selection. BOLD-fMRI images were used to localize the ROI within the visual cortex. The response map between 30-80 ms demonstrated a significant response in the visual cortex, as early as 40 ms after stimulus onset. **(B)** The participant underwent three conditions where the stimuli were presented 50 ms (stimuli onset 1), 40 ms (stimuli onset 2) and 30 ms (stimuli onset 3) prior to the scanning window. Average signal changes in the visual ROI revealed noticable time shifts across three conditions, corresponding to the different onset timings. The two markers indicate the time of peak 1 and peak 2, respectively.

Subsequently, we proceeded with an experiment comprising eight interleaved runs in which the stimuli were presented either 50 ms (stimuli onset 1) or 30 ms (stimuli onset 2) prior to the scanning window in a discovery group (N = 9). Each participant’s total acquisition time was approximately one hour. The group-average signals also exhibited a temporal shift in TRIGGER responses between these two conditions, with the signals of the onset 1 condition showing an earlier response compared to the 20-ms-delay signals (Fig. 3A). The group-average signals were not as sharp as the individual ones due to individual variability in response latencies. We then replicated the experiment in an independent group of 9 additional participants. The group-average signals again displayed a temporal shift in the responses between the two conditions (Fig. 3B), indicating that the approach can reproducibly captured the time differences in brain responses corresponding to a delay in the onset of stimuli.

**Fig. 3.**
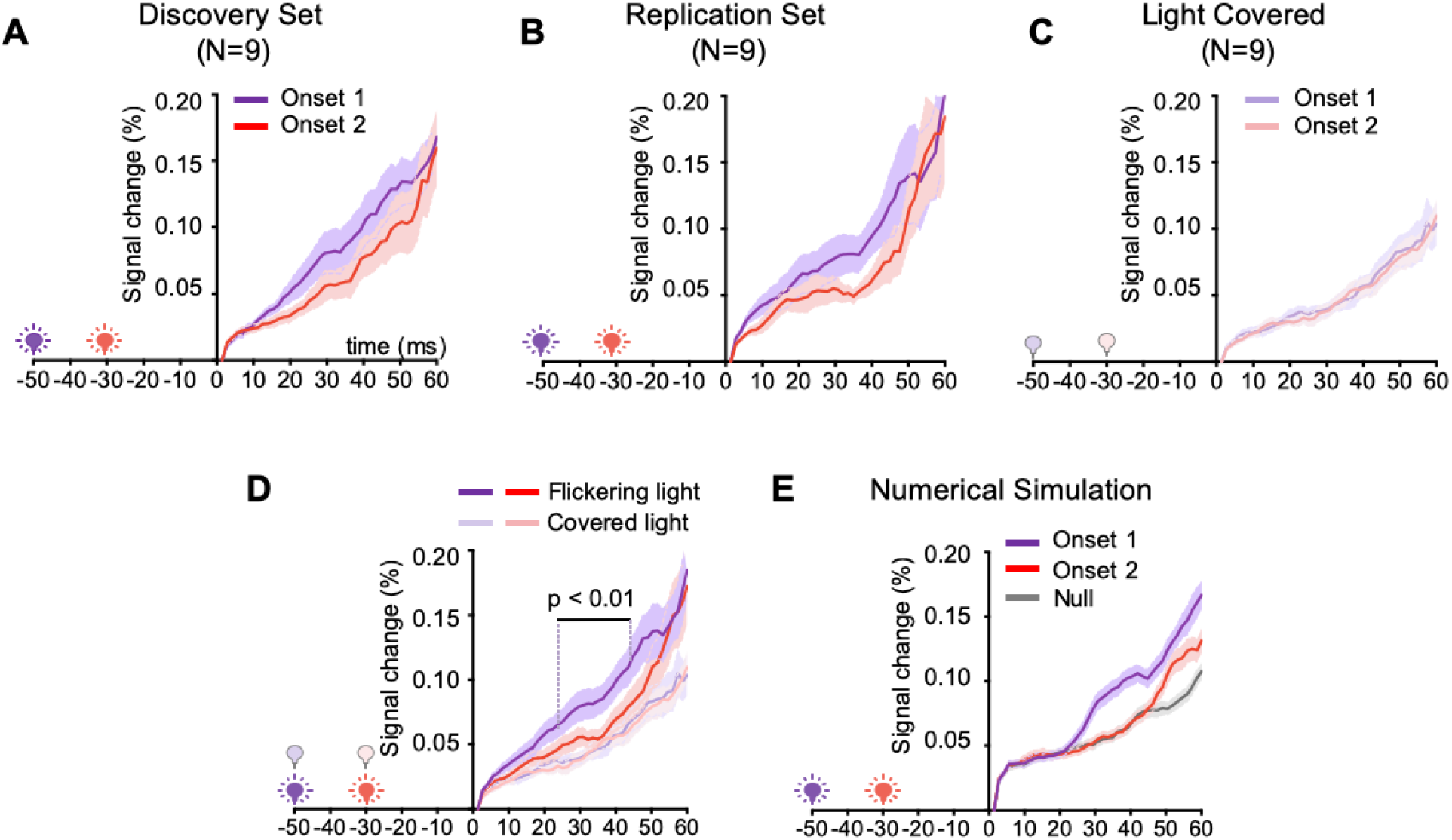
Reproducible millisecond-level responses captured by the TRIGGER at the group level. **(A)** An experiment with stimuli presented 50 ms (stimuli onset 1) and 30 ms (stimuli onset 2) prior to TRIGGER image acquisition window was conducted in a discovery group of 9 participants. Group-level signals revealed a time shift between two conditions, with an earlier response observed in the stimuli onset 1 condition compared to the stimuli onset 2 condition. (B) The experiment was replicated in an independent group of 9 participants. (C) A control experiment was conducted with the identical experimental settings in 9 participants, but with the flickering light completely covered. The group-level signals showed no difference between the two stimuli onset conditions. (D) Statistical analysis revealed significantly greater responses in the original condition, starting from 73.8 ms after stimulus onset, when comparing the flickering light setting (N=18) to the covered light setting (N=9). (E) Numerical simulation of signal changes in T2* of the two onsets and light covered conditions exhibited consistent patterns with the empirical data.

To rule out the possibility that the observed signal changes were actually artifacts stemming from the MRI sequence or the stimuli presentation system, we repeated the experiment in 9 participants with the identical experimental settings except that the flickering light was covered to prevent participants from receiving visual stimulation during the experiment. No significant response delay was found between the two stimuli onset conditions when the light was covered (Wilcoxon signed-rank test, all *p’s* > 0.05, Fig. 3C), despite a monotonous increase in signal changes which was expected. Our numerical simulation results indicated that the signal changes would still gradually increase when there is no biological activities (Fig. 3E, see Methods and Materials for details). Responses in the original stimuli condition were significantly greater compared to the stimuli-covered condition from 73.8 ms after stimuli onsets (Wilcoxon rank-sum tests, *p’s* < 0.01). These signals were highly consistent with results of the numerical simulations based on the canonical visual evoked potential (Fig. 3E). The post-stimulus latency of the TRIGGER responses appeared to be in line with that of the N75 component, the negative-going visual evoked potential component derived from EEG data (*16*).

Next, we performed an experiment in 10 participants to investigate the capacity of the TRIGGER approach to detect and differentiate the stimulus-evoked spatiotemporal dynamics within the left and right visual cortices. Each participant underwent ten interleaved scanning runs, during which visual stimuli were presented specifically to either the left or right visual field. Throughout the scanning sessions, participants were instructed to maintain fixation on a crosshair to ensure consistent gaze during the experiment (Fig. 4A). Stimuli presented to only one visual field are expected to evoke neural responses in the contralateral visual cortices earlier than the ipsilateral cortex (Fig. 4B). Indeed, when stimuli were presented to the left visual field, an earlier TRIGGER responses was observed in the right visual cortices compared to the left visual cortex (Fig. 4C). The pattern was reversed when the stimuli were presented to the right visual field (Fig.4D). We then combined these two results by averaging signal changes in the contralateral and ipsilateral visual cortices across all runs for each participant. We found significantly greater responses in the contralateral visual cortex compared to the ipsilateral visual cortex starting from 76.2 ms after the stimulus onsets (Fig. 4E, Wilcoxon signed-rank tests, *p’s* < 0.01)

**Fig. 4.**
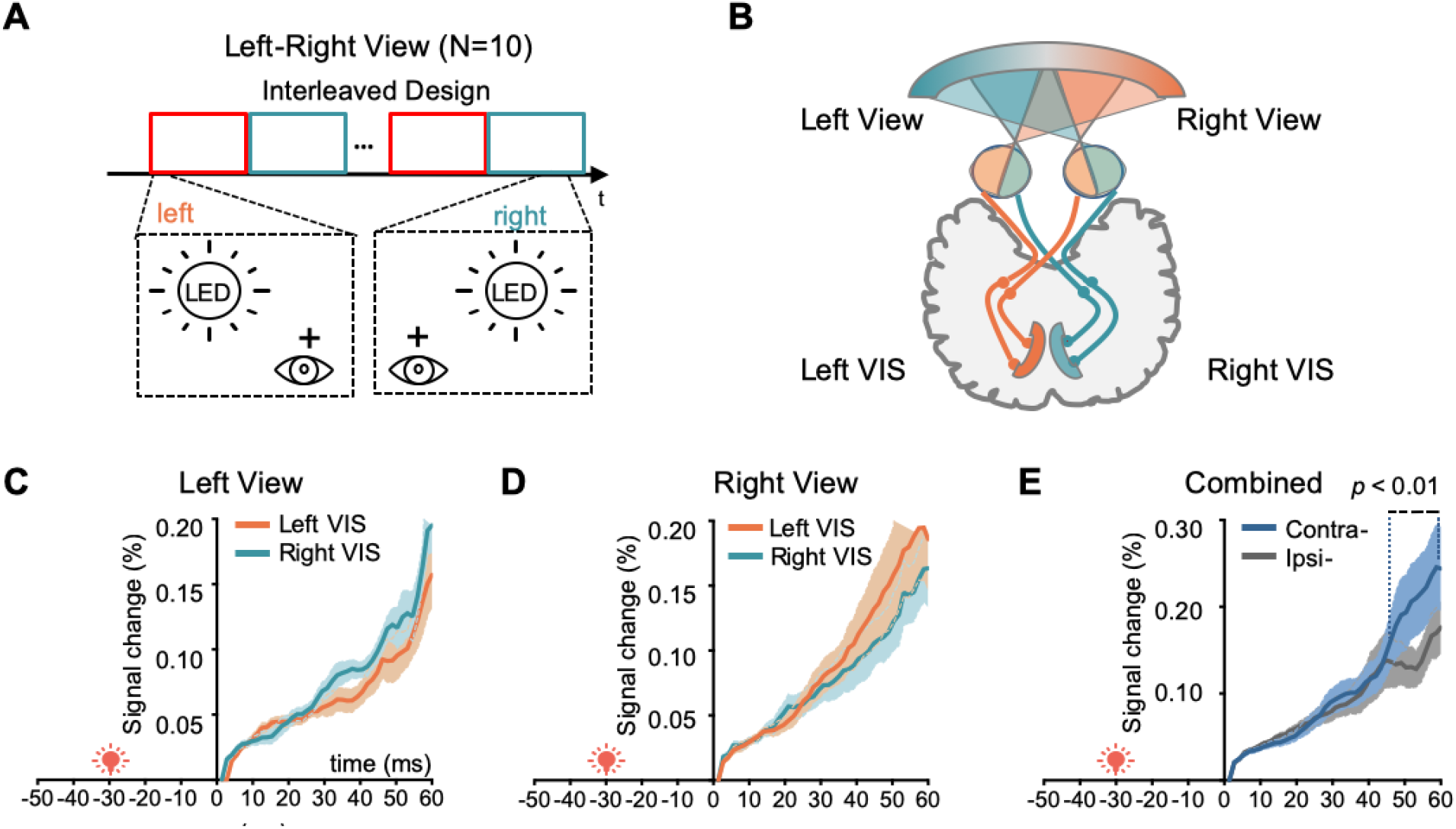
Dissociable spatiotemporal dynamics in the contralateral and ipsilateral early visual cortices. (**A**) Visual stimuli were presented to either the left or right visual fields of 10 participants, denoted as the Left View and Right View conditions, respectively. (**B**) A diagram of the primary pathway for visual signals from the retina to the visual cortex. VIS: early visual cortex. (**C**) When stimuli were presented in the left visual field, TRIGGER responses in the right visual cortex appeared earlier than those in the left visual cortex. (**D**) Conversely, in the Right View condition, TRIGGER responses in the left visual cortex appeared earlier than those in the right visual cortex. (E) Overall, TRIGGER responses within the contralateral visual cortex appeared earlier than those within the ipsilateral visual cortex. Statistical comparison reveals greater signal changes in the contralateral visual cortex starting from 76.2ms after the stimulus onset.

Finally, we performed a numerical simulation to examine whether the TRIGGER responses were dominated by the BOLD effect. The simulation incorporated a convetional hemodynamic response function with an initial dip to simulate the BOLD-fMRI responses with a 1% signal change, while a triangular waveform was used to emulate the TRIGGER responses with a 0.2% signal change (Fig. S1A). The simulated signals and the empirical BOLD-fMRI data showed similar dynamic patterns (Fig. S1B), suggesting that the simulation appropriately accounted for the BOLD effect. Using the identical analytical procedures as used for our experimental data, we were able to detect TRIGGER responses from the simulated signals (Fig. S1C).

## Discussion

In summary, the TRIGGER approach demonstrates the feasibility of capturing neural activity in awake humans with both high spatial and temporal resolutions using 3T MRI. Leveraging the high SNR offered by TRIGGER, we successfully localized the early visual cortices and discerned distinct responses to visual stimuli with varying onset timings on the order of milliseconds. Furthermore, by presenting stimuli to one visual field, we were able to distinguish spatiotemporal dynamics within the ipsilateral and contralateral visual cortices, highlighting the capability of TRIGGER to uncover intricate neural processes.

Translating the previously reported DIANA approach from anesthetized animals to awake humans proved to be challenging. One challenge arises from the subtle signal changes (∼0.1%), which require high SNR of MRI sequences for detection. While higher magnetic field strengths proportionally yield higher SNR, the safety of human participants limits scanner strength to commonly-used 3T or up to 10.5T (*17*). Another strategy to improve SNR is increasing the number of excitations (NEX), as was done in the DIANA study with the NEX of 40, but this leads to longer acquisition times. However, due to difficulty for participants in maintaining strict head fixation for longer periods, the duration of MRI sessions is limited (about 1.5 hours per participant in the study), making it challenging to increase NEX to improve SNR for human studies. These challenges led us to develop the novel MRI sequence to enhance SNR. Our event-synchronized echo-train-based sequence improved SNR during the time window of interest, which enabled us to obtain replicable TRIGGER signals evoked by visual stimuli in awake humans. Importantly, our signals captured the temporal shifts of stimuli onsets and may reflect spatiotemporal dynamics in the visual cortices of two hemispheres, suggesting its potential in recording neural activities in human studies. Our numerical simulations suggested that it is possible to detect TRIGGER responses from the background hemodynamic responses typically measured by BOLD-fMRI. Although we cannot entirely rule out the possibility that there is an undiscovered rapid hemodynamic response, the reproduciable and fast TRIGGER response to the visual stimuli is valuable and holds significant promise in human brain research.

The human brain is functionally organized into dynamic networks consisting of multiple adjacent and distant brain areas (*18-21*). However, understanding the intricate communication and interactions among these areas, as well as their connections with other functional networks, at a detailed level of resolution continues to pose challenges with the current available techniques. The TRIGGER provides high temporal resolution while simultaneously scanning multiple cortical and subcortical areas, offering a tool to mapping dynamic networks at a fine-grained scale. The technique may be particularly useful for studying higher-order cognitive circuits in humans, such as language, reasoning, and executive controls that involve long-range interactions. Moreover, by combining this high-resolution mapping technique with brain stimulation techniques one can explore the causal relationships between brain regions, leading to the formation of a brain causal connectome (*22*). Such combinations may also deepen our understanding of the neural mechanisms of brain stimulation techniques, such as deep brain stimulation (*23*) and transcranial magnetic stimulation (*24*), and lead to optimized stimulation parameters and targets to treat various brain disorders (*25, 26*).

Several important caveats should be taken into account when interpreting the results of this study. First, our current TRIGGER approach is preliminary and susceptible to the head motion. Participants need to stay still during the whole acquisition time, and thus the scanning cannot last a long time (up to 1.5 hours in awake human in our study). The acquisition time may be shortened in the future and the level of SNR could be maintained by reducing the NEX and the field of view, as well as adapting the parallel imaging strategy in ultrahigh-field MRI scanners. Second, due to the decays of SNR, the data acquisition window with sufficient SNR is relatively short (60 ms in the present study) thus the current TRIGGER approach can only sample part of the neural response. Third, both TRIGGER and DIANA may share the same imaging mechanism. Toi and colleagues previously hypothesized that these signal changes may arise from changes in membrane potential or cell swelling (*12*). However, further investigations in animals and patients are in needed to elucidate the exact origin of the signals.

In conclusion, we demonstrated the TRIGGER approach that can capture reliable signals in awake humans using 3T MRI. The potential to record neural responses in awake humans with high spatiotemporal resolution opens up avenues for uncovering dynamic functional organizations within the human brain and exploring their dynamic characteristics in cognitive processing. This advancement has the potential to drive significant discoveries in the fields of neuroscience and brain medicine as well as enhance our understanding of the complexities of the human brain.

## Materials and Methods

### Participants

We enrolled 18 young healthy participants (27.5 ± 2.8 years old, 11 women, 7 men). Prior to their inclusion in the study, we obtained informed consent from all participants, and they were screened for MRI contraindications. All participants had normal vision or corrected to normal vision before each scanning session. The Local Ethics Committee at the Changping Laboratory approved the study.

### MRI Experiments

All MRI experiments were performed in a 3.0-Tesla GE SIGNA UHP scanner (General Electric Healthcare, Waukesha, Wisconsin, US) equipped with a 48-channel head coil.

#### MRI experimental environment setup

To avoid introducing non-negligibly large and unstable latencies of stimuli after MRI acquisition onsets, we established a real-time stimuli presentation system to deliver visual stimuli. We elicited a synchronizing pulse signal from the MRI spectrometer to a STC89C51 microcontroller (STCmicro, Beijing, China), and the microcontroller controlled light-emitting diodes (LEDs) flickering (Fig. 1C & Fig. S2B). By measuring the time interval between the pulse signals of the MRI spectrometer and the optical signals of LEDs using an oscilloscope, we found that the latency of this system was 0 ms, when the measurement accuracy of oscilloscope is 1 ms. The latency of the system thus is less than 1 ms.

#### Anatomical sequence parameters

We performed an anatomical MRI scan to identify anatomical landmarks using the T1 sequence. The anatomical data had a TE of 24 ms, TR of 2000 ms, TI of 694 ms, FOV of 32 cm × 16 cm, acquisition matrix of 128 × 64, and five 5-mm coronal slices.

#### BOLD-fMRI sequence parameters

To localize the early visual cortices, we performed a BOLD-fMRI scan before the TRIGGER scanning. BOLD-fMRI images were acquired using a gradient-echo echo-planar imaging (EPI) sequence (TR = 3000 ms, TE = 30 ms, flip angle = 90°, FOV = 32 cm × 32 cm, acquisition matrix = 64 × 64, slice thickness = 5 mm, 5 coronal slices).

#### TRIGGER sequence parameters

To investigate neural activity with high spatiotemporal resolutions and high SNR with MRI, we implemented a periodic event encoded dynamic imaging sequence based on a gradient echo-train (*15*). In the sequence, each TR is synchronized with a cyclic event (e.g., visual stimuli onset) so that each echo in the gradient echo train can be positioned in a distinct k-space matrix, and all echoes in the echo train spread across a series of time-resolved k-space matrices (k-space 1, k-space 2, … k-space N). This process is repeated M times, each with a different phase-encoding step, until all 2D k-space matrices are adequately sampled. After applying 2D Fourier transforms to the k-space matrices, a set of time-resolved images are produced to characterize the underlying dynamic process. The temporal resolution in the sequence is determined by echo spacing (esp), which is 1.4 ms. The key parameters were TR = 300 ms, flip angle = 40°, esp = 1.4 ms, FOV = 32 cm × 16 cm, acquisition matrix = 64 × 32, slice thickness = 5 mm, NEX = 15, the number of discared dummy acquisitions = 20, and acquisition time = 5 minutes (500 trials) per run. Using the BOLD-fMRI retinotopic localizer task, we selected the slice with the most responsive voxels out of the 5 coronal slices for the TRIGGER scan.

#### DIANA sequence parameters

To compare the SNR between TRIGGER and DIANA, we implemented the DIANA sequence by adjusting a 2D gradient-echo imaging sequence, closely following the previous report (*12*). The key parameters were TR = 5 ms, TE = 2 ms, flip angle = 4°, FOV = 32 cm × 32 cm, acquisition matrix = 64 × 64, and slice thickness = 5 mm; To appropriately evaluate SNR, we used the NEX of 1 and the same FOV for both sequences, as well as the same acquired coronal slice for each participant.

#### Stimuli of BOLD-fMRI localizer

A block-designed retinotopic mapping experiment was conducted to localize the most responsive voxels in early visual cortex (*27, 28*). Specifically, we used the Sinorad stimuli presentation system (Sinorad, Shenzhen, China) to present a 45-degree wedge of contrast-reversing flicker, with a radius of 4.2°, a spatial frequency of 0.7 cycle/°, and a Michelson contrast of 1, against a uniform gray background with a mean luminance of 12.5 cd/m^2^. The wedge was presented on the upper, lower, left, or right visual field, with a contrast reversal rate of 6 Hz for 12 seconds, and the order of blocks was randomized. Participants were instructed to fixate on the central fixation point and remain still throughout the experiment.

#### Stimuli of TRIGGER visual experiments

To activate the early visual cortex in the TRIGGER experiments, we used LEDs to present flickering visual stimuli. Three LEDs, arranged in an inverted equilateral triangle with a side length of 2.7° and covered by a circle lampshade with a diameter of 5.7°, were placed on the upper visual field and repeatedly illuminated for 100 ms every 600 ms. In the pilot experiment and the experiment with varying stimuli onsets, participants were instructed to fixate at the central fixation, and LEDs located directly above with a visual angle of 3.4°. In the experiment to map left and right visual cortices, participants were instructed to fixate on a crosshair with visual angles of 5.7°. Participants were asked to remain still throughout the experiment.

### BOLD-fMRI data analyses

The BOLD-fMRI image preprocessing and analyses were performed using SPM12 (Wellcome Trust Center for Neuroimaging, Institute of Neurology, UCL, UK). Imaging preprocessing includes a spatial smoothing using a Gaussian kernel of 5-mm full width at half maximum (FWHM) and task activations were estimated using a general linear model. According to the task activations of the localizer task, we identified a ROI of the early visual cortex through including voxels with a response greater than 60% of the maximal response in the varying onsets experiment. In the Left-Right View experiment, the threshold of 60% was applied to left and right hemispheres, respectively.

### The TRIGGER data analyses

The TRIGGER signals were processed using the in-house MATLAB 2020b (MathWorks, Inc) codes with the following steps. Firstly, the timeseries were normalized through dividing by the first frame. Secondly, the absolute percent signal changes (PSC) were calculated using the formula, |TS_s_ – TS_0_|/TS_0_×100%, where TS_s_ and TS_0_ represent the signal changes after stimulation and baseline, respectively. Subsequently, temporal smoothing was applied using a centered moving average across 5 frames (7 ms). Furthermore, to map the TRIGGER responses in the brains, we performed permutation tests to examine the significant responses for each voxel at each timepoint. Specifically, we randomly selected 10 voxels as a random ROI within the cerebral mask and averaged timeseries within the random ROI. Random ROI selections were repeated 500 times to generate distributions of average signal-change timeseries. The signal change of each voxel was compared with the distribution of signal change and the comparison was calculated as a Z-statistic [(PSC_voxel_-mean of PSC_distribution_))/corrected standard deviation of PSC_distribution_] at each timepoint. The corrected standard deviation of PSC_distribution_ was the standard deviation of PSC_distribution_ multiplying by the square root of 10 to account for the lower noise level in 10-voxel ROIs compared to that in each single voxel. To control false positives from the multiple tests, when generating the significant response map, we excluded the voxels who either failed the statistical tests at both consecutive time points or had no neighboring voxel passing the tests.

### Numerical simulation analysis for SNR

We simulated the SNR of both the DIANA and TRIGGER sequence based on the following formula:

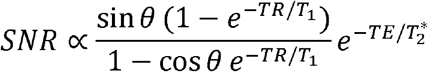

where *θ* represents the flip angle. To simulate the SNR of the DIANA sequence, we set the flip angle of 4° and the *TE* of 2 ms, according to the previous report (*12*). To simulate the SNR of the TRIGGER sequence, we set the flip angle of 40° and the *TE* ranging from 1.4 ms to 140 ms. The 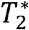 and *T*_1_ was set as 30 ms and 1100 ms for both sequences, respectively.

### Numerical simulation analysis for signal changes

According to the hypothesis derived from the DIANA study, alterations in membrane potentials or cellular swelling resulting from neural activities may elicit changes in T2* relaxation time. Considering the susceptibility of our imaging sequence to T2* variations, we postulated that the observed signal changes could be attributed to neural activity-induced T2* alterations. To investigate the hypothesis, a simulation analysis was conducted. Initially, we assumed a T2* value of 40 ms in the gray matter, a T2* variation amplitude of 0.4%, and the latencies of neural activity of 75 ms and 100 ms, with individual variability of 7 ms, consistent with the latencies of the event-related potential components N75 and P100 (*16*). Neurophysiological signals were simulated using a triangular waveform with a FWHM of 10 ms. Subsequently, the baseline FID was simulated using exp(-t/T2*) with an initial SNR of 5000. The stimulation FID was obtained by adding the 70-ms and 100-ms triangular waveforms to the baseline FID, mimicking the flickering light condition. No additional waveforms were added for the FID simulation under the light-covered condition. Finally, the same processing procedures with the empirical data were applied to generate simulated signal changes, and the simulation was repeated 100 times.

### Numerical simulation analysis for the BOLD effect

To investigate the impact of the BOLD effects on TRIGGER responses, a 300-second signal was simulated to incorporate both the BOLD and TRIGGER components. Initially, we assumed that each stimulus with an ISI of 600 ms would elicit a hemodynamic response with a 1% signal change and a corresponding TRIGGER response with a 0.2% signal change. To simulate the BOLD effect, an impulse response function with varying amplitudes randomly sampled from a Gaussian distribution (mean=1, standard deviation=0.5) was convolved with a canonical hemodynamic response function that accounted for an initial dip (*29*). For the TRIGGER response simulation, a triangle waveform with a post-stimulous latency of 80 was utilized. Subsequently, the simulated BOLD signals, TRIGGER signals, and white noises with a SNR of 300 were combined to obtain the simulated response signals. Finally, the same processing procedure with the empirical data were applied to estimate the simulated signal changes.

### Statistical analyses

Wilcoxon rank-sum tests were used to compare the signal change at each timepoint under the flickering and covered light conditions. Wilcoxon signed-rank test was used to compare the signal change in contra- and ipsi-lateral visual cortices.

---

**Fig. S1.**
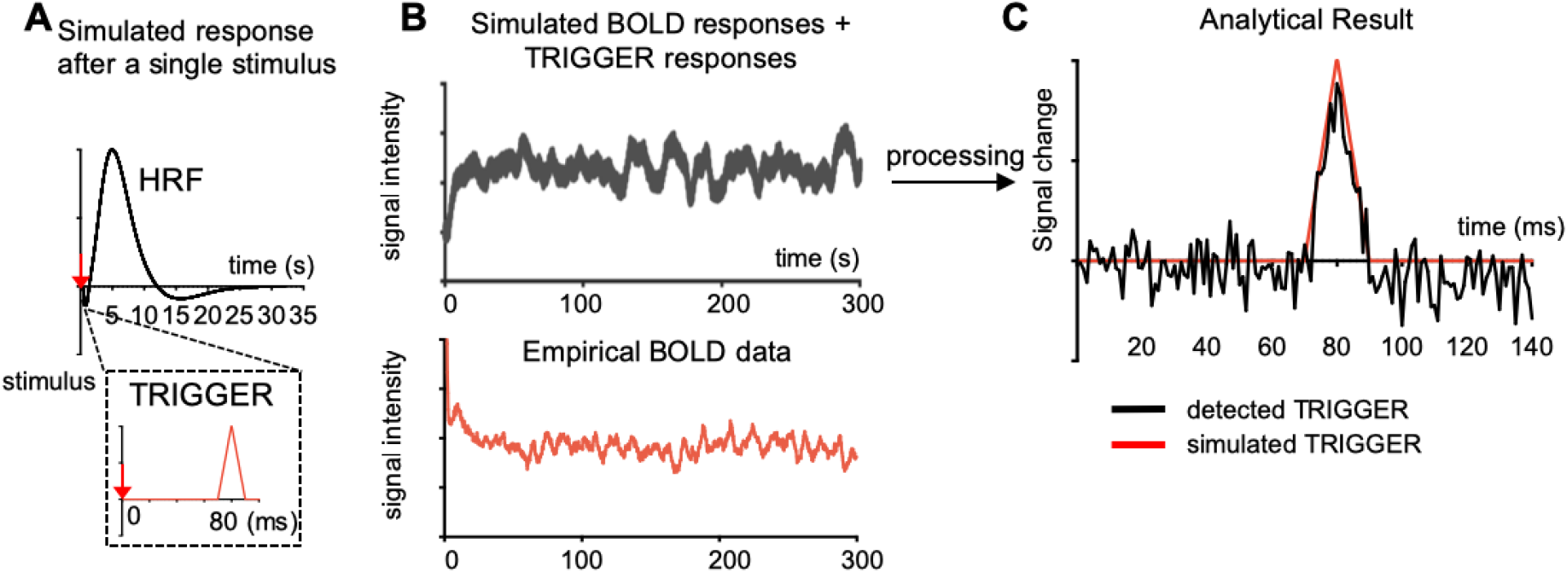
The numerical simulating analysis of the impact of BOLD effect on the TRIGGER responses. (A) A simulated BOLD response and a TRIGGER response were generated in response to a single stimulus. The BOLD response was simulated using a hemodynamic response function, while the TRIGGER response was simulated using a triangle waveform with a post-stimulus latency of 80 ms. (B) Simulated signals incorporating both BOLD and TRIGGER responses were generated over a duration of 300 seconds under a condition that stimuli repeated at an ISI of 600 ms. The amplitudes and fluctuations are similar to those observed in the empirical BOLD data. (C) The TRIGGER responses could be detected from the simulated signal through the identical analytical procedure as used for our experimental data.

## Notes

### Competing Interest Statement

The authors have declared no competing interest.

